# Deciphering the role of the OmpC-Mla system in bile salt resistance

**DOI:** 10.1101/2023.06.27.546672

**Authors:** Feifan Zhu, Zhi-Soon Chong, Shu-Sin Chng

**Affiliations:** Department of Chemistry, National University of Singapore, Singapore 117543; Singapore Center for Environmental Life Sciences Engineering, National University of Singapore (SCELSE-NUS), Singapore 117456

**Keywords:** Outer membrane, OmpC-Mla, MlaD, bile salt transport, deoxycholate, lipid binding

## Abstract

The outer membrane (OM) of Gram-negative bacteria presents a formidable barrier against external insults, in part contributing to the survival of Enterobacteriaceae in the mammalian gut. The lipid asymmetry of the OM, where lipopolysaccharides (LPS) form a tight outer layer of low permeability, effectively restricts the passage of toxic substances across the bilayer. In the gut, however, bile salts pose a unique challenge to the bacterial cell due to their ability to form micelles and solubilize membranes; yet, mechanisms to prevent dissolution of the OM by such detergents are not well understood. In this study, we define a distinct role in bile salt resistance for the OmpC-Mla system in *Escherichia coli*, which is better known for its function in maintaining OM lipid asymmetry. We show that cells lacking a functional OmpC-Mla system are sensitive to bile salts, but only at or above critical micellar concentrations. Furthermore, we observe that these cells still exhibit bile salt sensitivity even when defects in OM lipid asymmetry have been corrected, suggesting that the OmpC-Mla system contributes to bile salt resistance independent of its role in lipid asymmetry. Finally, we demonstrate that MlaD, one of the key lipid-binding components of the system, displays specific binding to bile salts in vitro. Since the OmpC-Mla system maintains OM lipid asymmetry by transporting mislocalized PLs, our findings here support a model where this system also additionally removes bile salts that have intercalated into the OM, to ultimately prevent dissolution and disruption of this important barrier.

**IMPORTANCE:** Bile salts are important components secreted into the human gut to help solubilize fats from our diet, yet they also possess anti-microbial properties due to their corresponding ability to dissolve bacterial membranes. For Enterobacteriaceae to survive in the gut environment, these bacterial cells must prevent intracellular build-up of bile salts by either restricting entry or by pumping out these molecules. Ultimately, they must resist bile salt-mediated dissolution of their membranes, particularly their outer membranes, which serve as a protective barrier against toxic substances. In this study, we reveal that a known lipid transport system in *Escherichia coli* has a distinct role in bile salt resistance independent of its role in maintaining outer membrane lipid asymmetry; it does so likely by removing bile salts from the outer membrane, thus preventing dissolution. Our work highlights the possibility of targeting this lipid transport system for the treatment of Enterobacteriaceae infections.

## INTRODUCTION

Enterobacteriaceae are a group of Gram-negative bacteria, including *Escherichia coli* and *Klebsiella pneumoniae*, that inhabit the gastrointestinal tracks of animals. In humans, these bacteria are commensals that play important roles in host metabolism, immune, and neuroendocrine responses [1], but can also become opportunistic pathogens, such as causing late-onset sepsis. Enterobacteriaceae need to survive harsh conditions to live in the mammalian gut. In particular, the gut contains large amounts of bile salts (average 0.415% by weight) [2], which act as detergents that can solubilize lipids to aid food digestion, and thus represent a key stressor on bacterial membranes. The major bile salts are typically cholate, chenodeoxycholate (CDOC), and deoxycholate (DOC), predominantly in glycine/taurine-conjugated versions.

To resist the action of bile salts, Gram-negative bacteria possess at least two known mechanisms to prevent build-up of these detergent molecules in their membranes. The distinct double-membrane cell envelope structure features an outer membrane (OM) that serves as a barrier against the entry of bile salts into the cell. The OM is asymmetric with lipopolysaccharides (LPS) in the outer leaflet and phospholipids (PLs) in the inner leaflet. The well-packed LPS layer possesses low fluidity and restricts both hydrophobic and large hydrophilic molecules from entry [3]. In their monomeric form, however, bile salts are believed to be able to permeate across the OM through diffusion porins, including OmpC and OmpF. To protect against these bile salt molecules accumulating in and dissolving the inner membrane (IM), Gram-negative bacteria also express efflux pumps to extrude such harmful substances. In *E. coli* specifically, AcrAB-TolC constitutes the major efflux pump, where the homotrimeric Resistance-Nodulation-Division family protein AcrB connects to the OM homotrimeric TolC pore via the membrane fusion protein AcrA to form a continuous conduit for expulsion of molecules from the periplasm and the IM [4]. Being detergents, bile salts also form micelles above their critical micelle concentrations (CMCs). In fact, given the typical concentrations of bile salts in the gut, these molecules likely exist almost exclusively in micelles. Bile salt micelles have the ability to partition into the OM bilayer itself, perhaps aided by specific interactions between bile salts and LPS [5]. Accumulation of bile salts in the OM will lead to dissolution. Since efflux pumps cannot access substrates at the OM, how cells potentially remove bile salts that could have partitioned into the OM is not clear. Sodium taurocholate is known to induce OM vesicle production in *Campylobacter jejuni*, providing a possible avenue for bile salt removal [6].

While OM porins are generally regarded as channels for diffusive entry of bile salt molecules, how they contribute to bile salt resistance remains elusive. Specifically, removing OmpC, but not OmpF, actually makes cells sensitive to bile salts [7-9]. Thus, the presence of OmpC is important for bile salt resistance, which is unlikely due to its role as a diffusion porin. Interestingly, both OmpC and OmpF have recently been shown to associate with the OM lipoprotein MlaA, which plays a role in maintaining OM lipid asymmetry as part of the Mla system [10]. Yet, only removing OmpC, but not OmpF, causes defects in lipid asymmetry, which ultimately impacts the OM permeability barrier. This correlation raises the question of whether the role of OmpC in bile salt resistance may be attributed to its function within the OmpC-Mla system.

The OmpC-Mla system comprises seven proteins that work together to maintain OM lipid asymmetry [10, 11]. It does so by mediating the ATP-dependent retrograde transport of PLs from the OM to the IM [12, 13]. The periplasmic chaperone, MlaC, is believed to extract PLs from the outer leaflet of the OM via the OmpC-MlaA complex, and hand them over to the MlaFEDB ATP-binding cassette (ABC) transporter at the IM. MlaD is a mammalian cell entry (MCE) domain-containing single-pass membrane protein; it forms a hexameric donut structure with a central hydrophobic pore in the periplasm, and associates with a dimer of MlaFEB [14, 15]. The MlaD hexamer has been shown to co-purify with up to four lipids [14], but structures of other MCE proteins suggest possibly six binding sites [16]. In the MlaFEDB complex, the hydrophobic pore of MlaD hexamers forms a continuous lipid binding cavity with the trans-membrane region of the MlaE dimer [12, 15, 17, 18].

Here, we sought to understand the possible involvement of the OmpC-Mla system in bile salt resistance. We show that removing any member of the system results in sensitivity against bile salts at or above CMCs. This is distinctly different from sensitivity conferred by the loss of the AcrAB-TolC efflux pump. We demonstrate that this contribution to bile salt resistance is independent of the established function of the OmpC-Mla system in maintaining OM lipid asymmetry. Interestingly, we find that MlaD is capable of specifically binding bile salts in vitro, suggesting that it may have alternative substrates. Given that the Mce4 homologous system in mycobacteria is believed to transport cholesterol [19], the structural precursor to bile salts, our work here points towards a plausible mechanism for Gram-negative bile salt resistance where the OmpC-Mla system removes bile salts that have partitioned into the OM to prevent dissolution.

## RESULTS

### The OmpC-Mla system is important for resistance against bile salts at or above CMCs

The trimeric OmpC porin has been implicated in bile salt resistance [8]. To test if the rest of the OmpC-Mla system contributes to this phenotype, we examined sensitivity of deletion mutants to DOC, CDOC, and cholate, which are the three major bile salts in the human gut. As previously reported [7], the deletion of *ompC*, but not *ompF*, conferred bile salt sensitivity (**Fig. 1A, S1A**). We found that individual *mla* mutants also exhibited sensitivity towards bile salts, and this phenotype could be complemented (**Fig. 1B, S1B**). Importantly, the OmpC_R92A_ variant also displayed similar bile salt sensitivity (**Fig. 1C, S1C**). This OmpC variant is still functional as a diffusion porin yet gives rise to OM lipid asymmetry defects [20], indicating that the role of OmpC in bile salt resistance is independent of its porin function but dependent on its function with the Mla system.

**Figure 1:**
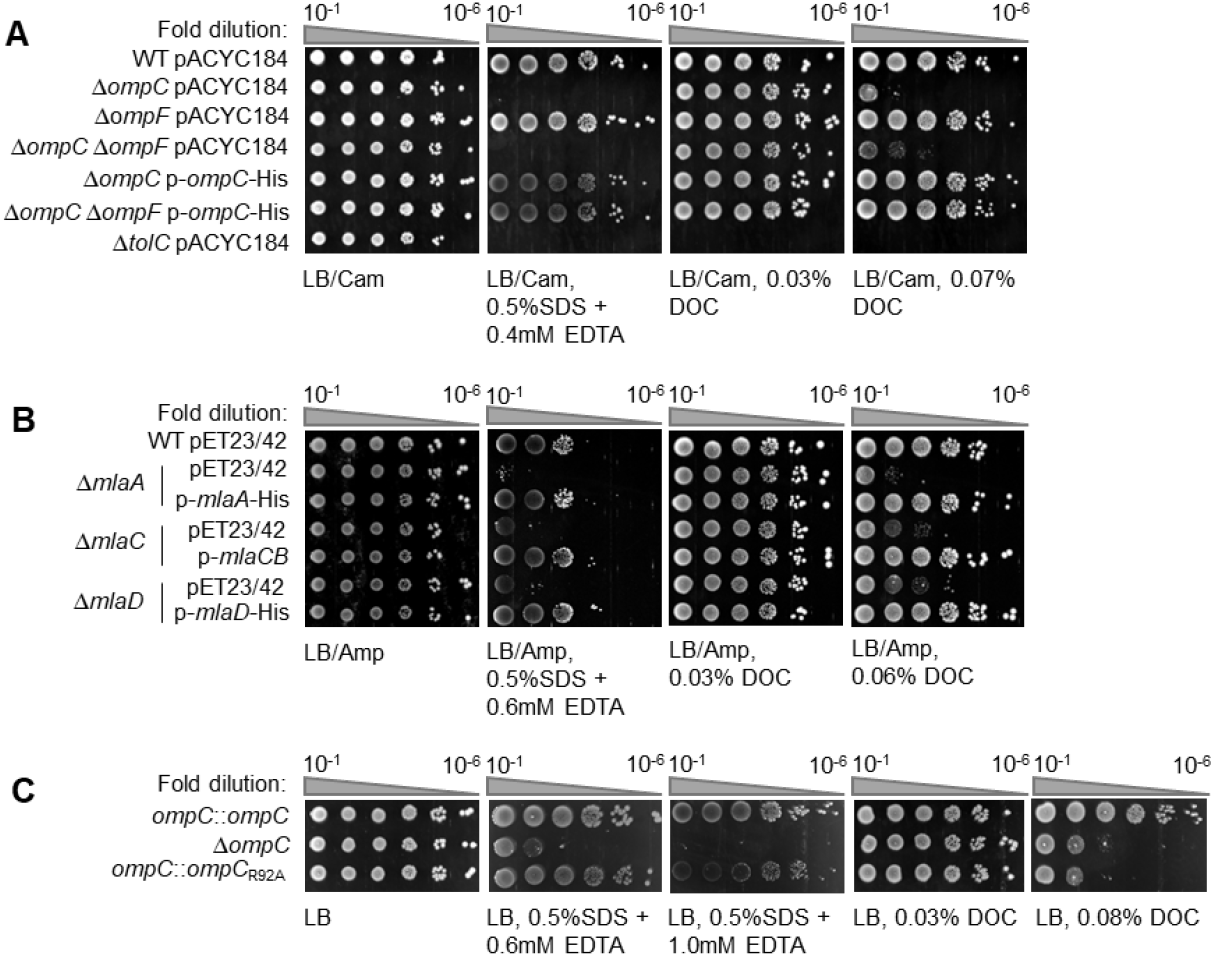
Defects in the OmpC-Mla system confer bile salt sensitivity at/above CMCs. Efficiency of plating (EOP) of indicated porin (A, C) or *mla* mutant (B) strains on LB media containing indicated concentrations of SDS/EDTA or DOC. *ompC-mla* strains exhibit sensitivity to SDS/EDTA at different minimum EDTA concentrations, likely due to varying Mg^2+^ content in different media batches [30]. Likewise, varying salt content could have impact on CMC values, and hence the minimum DOC concentrations that confer sensitivity. CMC values for DOC in buffers containing 0.1 M NaCl are in the range of 0.095-0.11% [22]; since LB is a complex media, we assumed that concentrations of DOC ≥ 0.06% in LB is at/above CMC.

We noted that *ompC-mla* mutants might only become sensitive around or above the corresponding CMCs of DOC, CDOC, and cholate (**Fig. 1**). The CMCs of these three bile salts in water (%w/v) are reported to be 0.10∼0.27%, 0.12∼0.26%, and 0.27∼0.55%, respectively [21]; in LB medium, these are expected to be lower due to increased ionic strength [22]. In contrast, deletion of *tolC* displayed bile salt hypersensitivity at concentrations likely lower than CMCs (**Fig. 1A, S1A**). We conclude that the OmpC-Mla system contributes to bile salt resistance at or above CMCs via a mechanism different from efflux systems.

### The OmpC-Mla system contributes to bile salt resistance beyond its role in maintaining OM lipid asymmetry

Cells lacking the OmpC-Mla system exhibit defects in lipid asymmetry, and thus permeability, in the OM. To understand if bile salt sensitivity in *ompC-mla* mutants is contributed solely by OM defects, we restored OM lipid asymmetry by overexpressing the OM phospholipase PldA to degrade PLs in the outer leaflet of the OM. Overexpression of PldA fully rescued SDS/EDTA sensitivity of ∆*mla* mutants (**Fig. 2A, S2A**), as previously reported [11]. However, doing so did not fully rescue bile salt sensitivity. We observed similar rescue (or lack of) in these phenotypes in *mla* mutants defective in ATP hydrolysis (*mlaF*_*K47R*_ and *mlaB*_*T52A*_) [14] (**Fig. S2B**). We further showed that PldA overproduction did not restore bile salt resistance in the *ompC*_*R92A*_ strain while completely rescuing OM asymmetry defects, and hence SDS/EDTA sensitivity (**Fig. 2B, 2C, 2D**). It appears that the OmpC-Mla system actively contributes to bile salt resistance in part independent of its role in OM lipid asymmetry.

**Figure 2:**
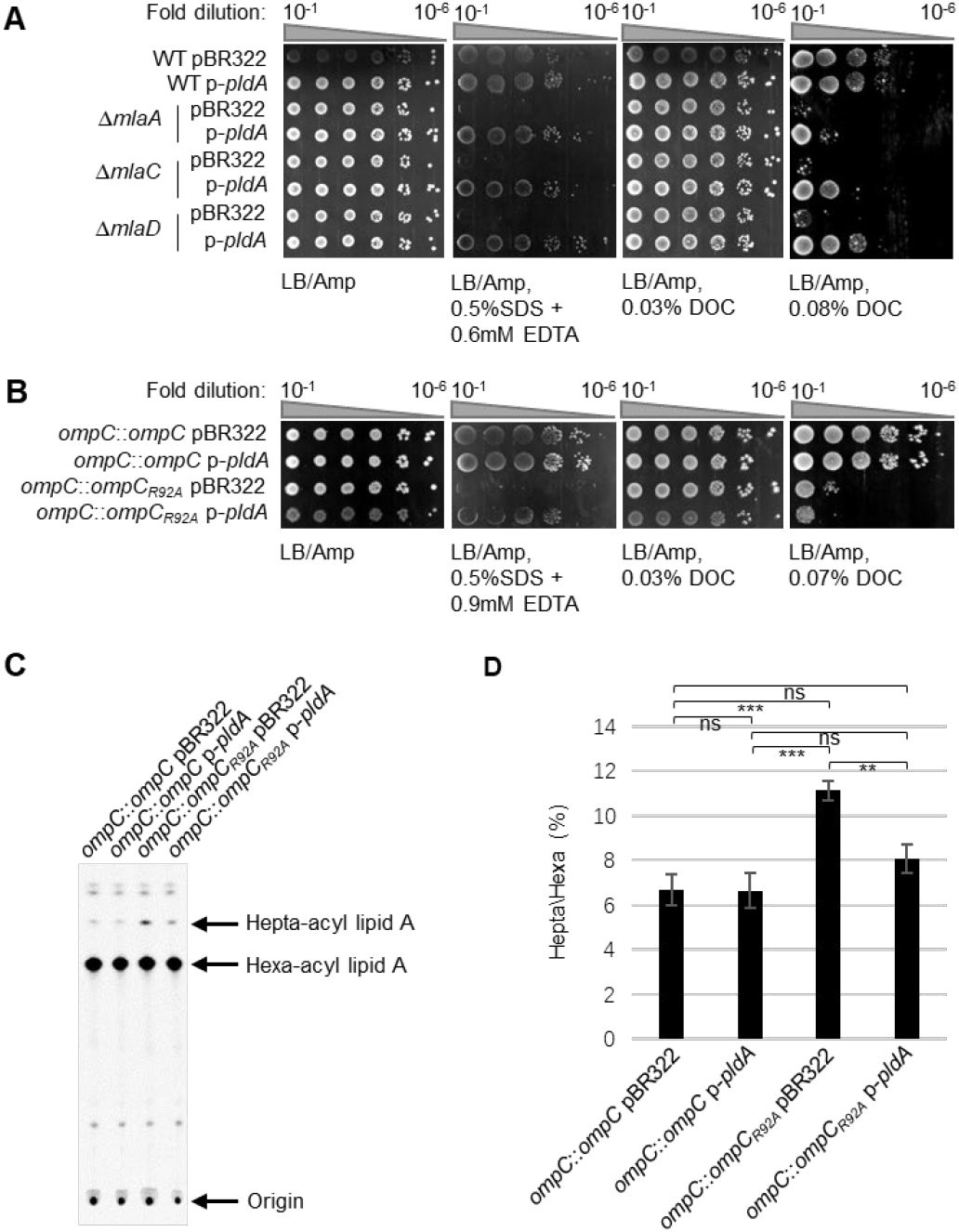
Overexpression of PldA fully rescues SDS/EDTA sensitivity but partially restores bile salt resistance in *mla* knockouts and the *ompC*_*R92A*_ mutant. EOP of *mla* mutant (A) or *ompC* mutant (B) strains harboring empty vector (pBR322) or p-*pldA* on LB media containing indicated concentrations of SDS/EDTA or DOC. (C) TLC/autoradiographic analyses of [^14^C]-labeled lipid A extracted from exponential phase cultures of *ompC* mutant strains harboring empty vector (pBR322) or p-*pldA*. (D) Average percentages of palmitoylation of lipid A from (C)and the standard deviations were quantified from triplicate experiments. Student’s t-tests: ns, not significant; **, p < 0.01; ***; p < 0.001.

### MlaD interacts specifically with bile salts in vitro

Since the OmpC-Mla system is a PL transporter, and homologs in mycobacteria such as the Mce1 and Mce4 systems are believed to transport fatty acids and cholesterol, respectively [19, 23], we hypothesize that the OmpC-Mla system may transport other hydrophobic substrates like bile salts. To test this idea, we examined if MlaD, the lipid-binding MCE domain protein, is capable of interacting with DOC. MlaD exists as a hexamer with a hydrophobic central pore connected to six putative substrate binding cavities at protomer interfaces [24], where lipid substrates from MlaC have been postulated to passage through to enter the MlaFEDB complex [24, 25]. We purified the soluble domain of MlaD (sMlaD) in apo- and holo-forms [24], and analyzed possible interactions with DOC using isothermal titration calorimetry (ITC). As a specificity control, we first demonstrated that DOC does not bind the soluble domain of LolB (sLolB), the OM lipoprotein receptor that contains a hydrophobic cavity (**Fig. 3A**). Remarkably, we found that apo-sMlaD displayed moderate-to-weak saturable binding to DOC, characterized by a sigmoidal titration profile; one hexamer binds six DOC molecules at K_D_ ∼100 μM (**Fig. 3B**). The same results were observed for CDOC (**Fig. 3C**). In contrast, apo-sMlaD only exhibited non-saturable binding to the octyl β-D-glucopyranoside (OG) detergent, indicating that its interactions with DOC and CDOC are specific (**Fig. 3D**). Importantly, holo-sMlaD, which contains four bound phospholipids [24], exhibited minimal interactions with DOC except during the first few titrations, possibly reflecting DOC binding in the MlaD central pore (**Fig. 3A**). We conclude that MlaD interacts with bile salt molecules in a specific manner likely involving use of its putative substrate binding pockets.

**Figure 3:**
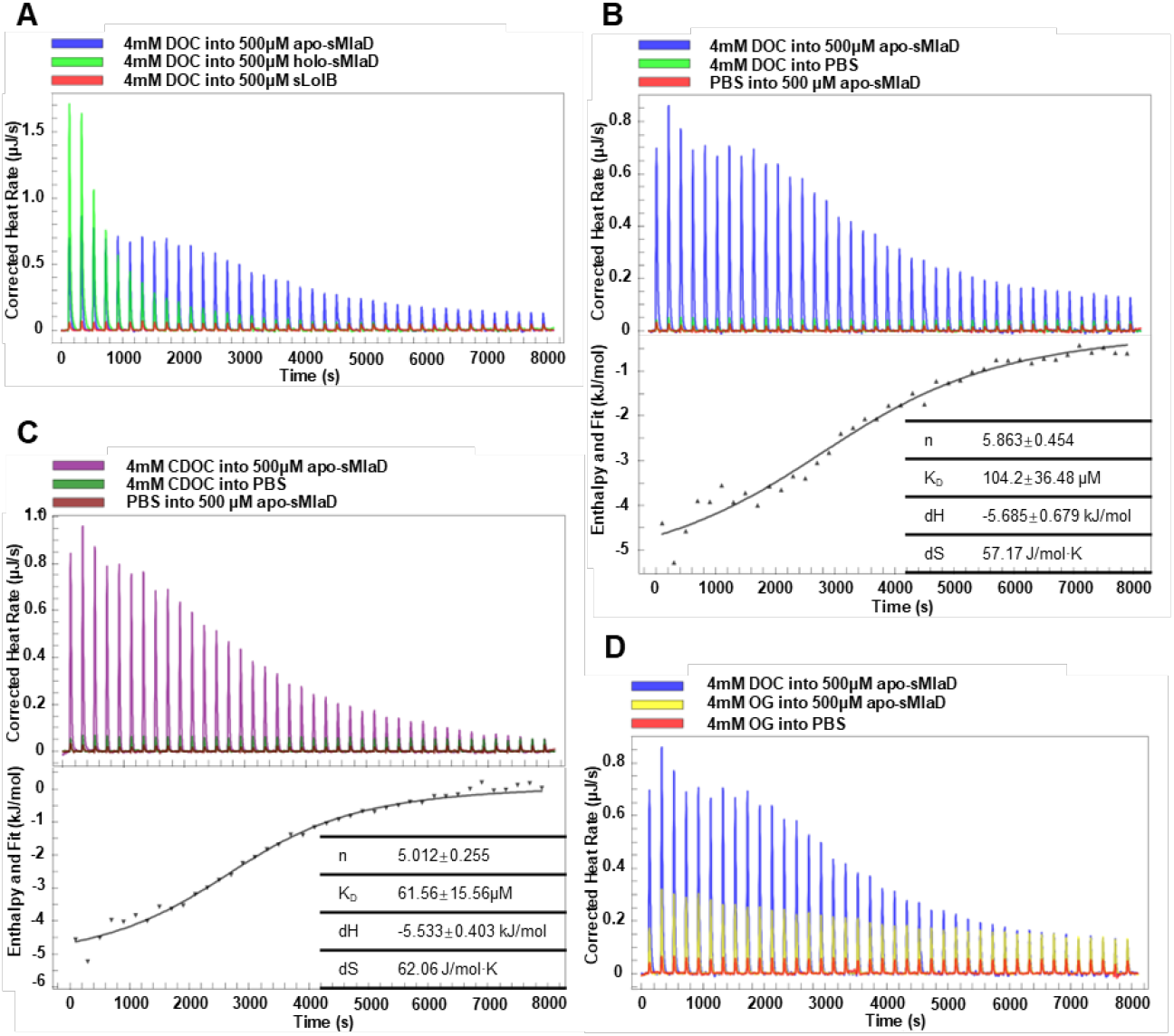
MlaD interacts specifically with bile salts in vitro. Representative ITC curves showing heat changes as DOC was titrated into (A) sLolB, holo-sMlaD, or (B) apo-sMlaD, and as (C) CDOC and (D) OG were titrated into apo-sMlaD. Titrations of DOC/CDOC into buffer (PBS), or buffer into protein solution are used as controls. For DOC/CDOC titrations into apo-sMlaD, curve fitting after background subtractions provided binding stoichiometry (n) and affinity (K_D_) estimates as shown.

We next applied photo-crosslinking to further validate binding specificity between DOC and apo-sMlaD. We incubated apo-sMlaD with a diazirine-containing DOC analog (az-DOC) and probed for covalent adduct formation upon UV irradiation (**Fig. 4A**). Consistent with our ITC data, az-DOC readily formed crosslink adducts with apo-sMlaD, but not sLolB (**Fig. 4B**). Furthermore, we showed that DOC, CDOC and cholate were all able to prevent the formation of az-DOC-apo-sMlaD crosslinks in a concentration-dependent manner (**Fig. 4C, 4D, S3**). To pinpoint potential site(s) of interaction, we subjected the az-DOC-apo-sMlaD adduct to trypsin digestion, and applied tandem mass spectrometry to sequence resulting DOC-crosslinked peptides. Interestingly, the residue D95, positioned close to the MlaD protomer interfaces, was identified as a high-confidence crosslink site (**Fig. S4A, S4B**). Taken together, our results establish saturable and competitive binding between DOC and apo-sMlaD, providing strong evidence for a specific interaction.

**Figure 4:**
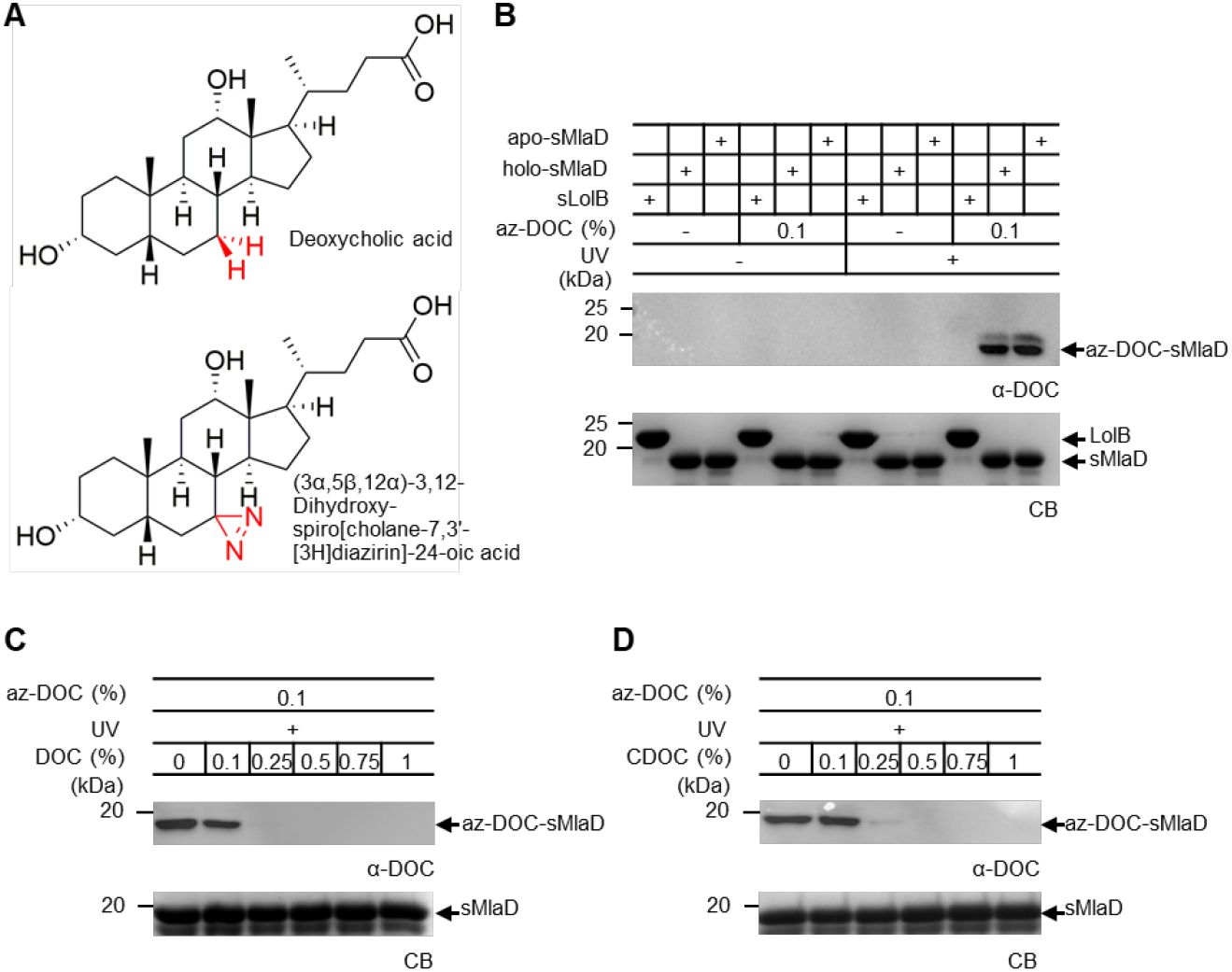
az-DOC crosslinks specifically to MlaD. (A) Chemical structures of DOC and (3α,5β,12α)-3,12-Dihydroxy-spiro[cholane-7,3’-[3H]diazirin]-24-oic acid (az-DOC). SDS-PAGE analyses and immunoblots showing UV-dependent formation of az-DOC crosslinks with (B) sMlaD, and in the presence of increasing concentrations of (C) DOC, or (D) CDOC. CB, Coomassie blue staining.

## DISCUSSION

It has long been appreciated that the presence of OmpC, but not OmpF, is required for bile salt resistance in Enterobacteriaceae [7, 8]. These trimeric porins are highly similar, yet why OmpC has a specific role in this phenotype is unclear. Here, we have uncovered that the requirement for OmpC in bile salt resistance is independent of its porin function, but instead dependent on its functional association with the Mla system [10]. Specifically, we have demonstrated that *ompC-mla* mutants exhibit bile salt sensitivity at/above CMCs (**Fig. 1**). In addition, we have shown that this sensitivity is not simply due to the resulting OM lipid asymmetry defects found in these strains (**Fig. 2**). We have also established that MlaD binds bile salt molecules in a specific manner in vitro (**Fig. 3, 4**), suggesting a possibility that the OmpC-Mla system may remove bile salts from the OM to confer resistance. Our work provides insights into how Enterobacteriaceae like *E. coli* resists high concentrations of bile salts in the mammalian gut.

We propose distinct mechanisms for how *E. coli* resists bile salts under two different scenarios, specifically below or above the CMCs (**Fig. 5**). At concentrations below CMCs, bile salts predominantly exist as monomers. As bile salt molecules cross the OM via diffusion porins and gain access to the IM, they can be pumped out by the AcrAB-TolC system. At concentrations above CMCs, bile salts form micelles, which can partition into the outer leaflet of the OM due to its solubilizing property, particularly its ability to disperse LPS [5]. Since the AcrAB-TolC system does not have access to substrates at the OM, the cell requires a mechanism that could remove these detergent-like molecules to prevent accumulation in and dissolution of the OM. Our results suggest that the OmpC-Mla system may provide this function. Given that this system extracts PLs from the outer leaflet of the OM to shuttle them back to the IM, and that MlaD interacts specifically with bile salts (**Fig. 3, 4**), it may be conceivable that the OmpC-Mla system could also remove other hydrophobic substrates (e.g. bile salts) from the outer leaflet of the OM, facilitating access by efflux pumps. Of course, one caveat of this model is that the observed interaction between MlaD and DOC in vitro appears to be weak, and may not be consistent with specific transport. Yet, it is known that when associated with MlaFEB, the lipid binding cavity within the MlaFEDB complex is significantly augmented by the transmembrane regions of MlaE [12, 15, 17, 18]. Affinity for lipids and/or bile salts in the native MlaFEDB complex may be sufficiently high to enable transport [13]. Consistent with the requirement for the full ABC transporter, bile salt resistance is compromised in mutants with defective ATP hydrolytic activity (**Fig. S2B**). The possible role and contribution of the IM MlaFEDB complex in transporting bile salts warrants further examination.

**Figure 5:**
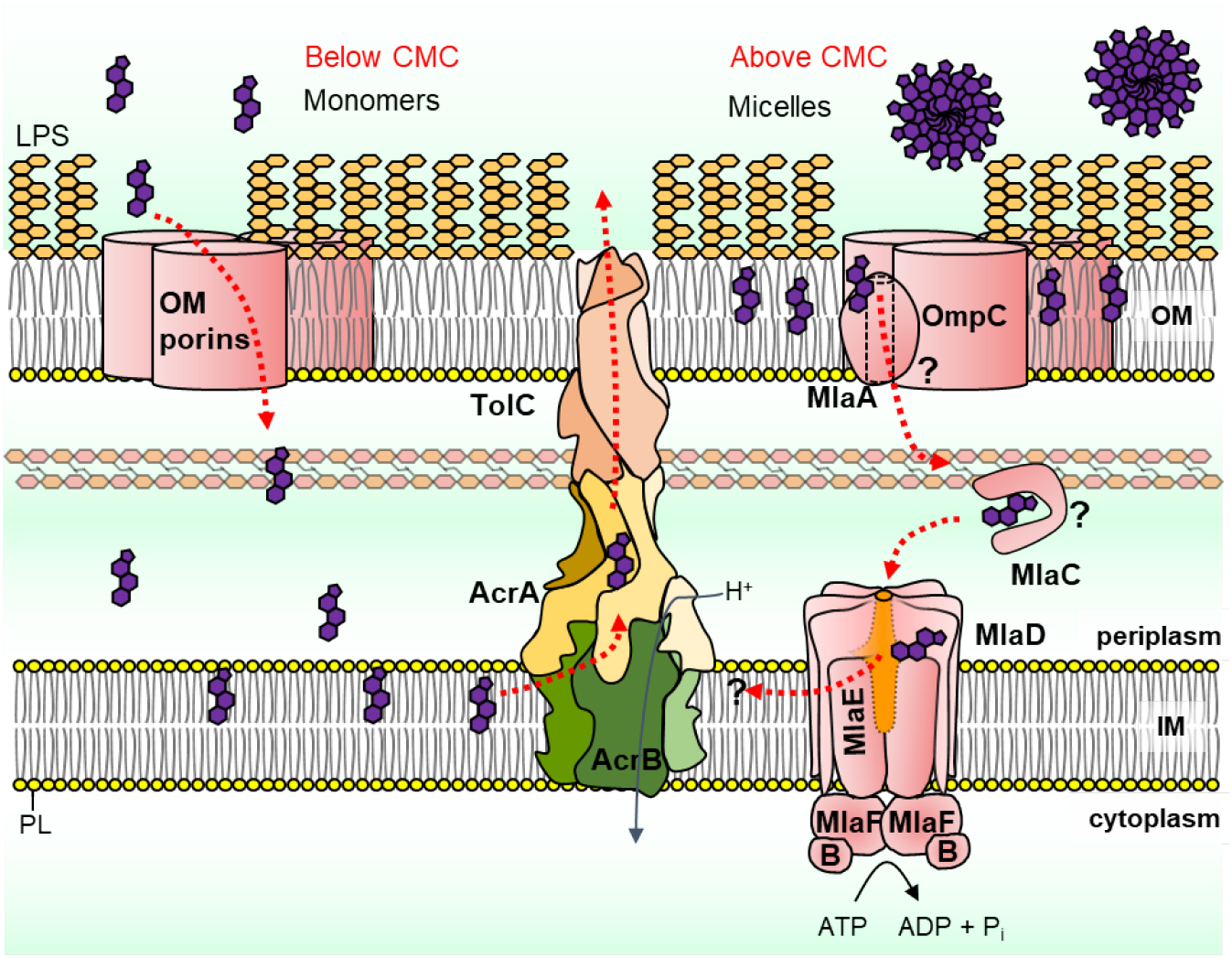
A proposed model for how the OmpC-Mla system contributes to bile salt resistance, particularly above CMCs.

There are two proteins, MlaC and MlaD, in the OmpC-Mla system that have been shown to bind lipids [14]. At this stage, we did not detect any binding between MlaC and DOC in vitro, either by ITC or crosslinking. However, we are confident that MlaC is required for bile salt resistance, as ∆*mlaC* mutant was still sensitive when we expressed *mlaB* to counteract the polar effect that *mlaC* deletion had on downstream *mlaB* (**Fig. S5**). If the OmpC-Mla system indeed transports bile salts from the OM, one possibility could be that the MlaC lipid binding pocket only opens upon interacting with partners such as MlaA or MlaD [26, 27]. Another possibility is that bile salt molecules may be co-transported with PLs in the event of accumulation in the OM. The lipid binding cavity of MlaC can accommodate binding of up to four acyl chains in *Pseudomonas putida* [26] and *Pseudomonas aeruginosa* [28]; such plasticity may allow MlaC to bind one PL molecule along with one other hydrophobic substrate (like bile salts). Additional studies would be required to validate these speculative ideas.

Our study reveals the novel role of the OmpC-Mla system in bile salt resistance, possibly explaining why OmpC, but not OmpF, is important for survival in high concentrations of bile salts in previous studies [7-9]. The Mla system belongs to the family of MCE systems believed to have roles in lipid transport. In *E. coli*, two other MCE systems include PqiABC and YebST (LetAB) [29]. The Pqi system has been shown to be required for resistance against the certain zwitterionic detergents like lauryl sulfobetaine. Since OmpC-Mla system contributes to bile salt resistance, the MCE family could possibly have unappreciated roles in removing general hydrophobic contaminants from the OM. Given that bile salt resistance is particularly important for Gram-negative bacterial colonization and survival in the mammalian gut, the OmpC-Mla system (and other MCE complexes) may be possible targets for future development of anti-infectives.

## EXPERIMENTAL

### Strains, plasmid construction, and growth conditions

All bacteria strains and plasmids used in this chapter are summarized in **Table S1** and **Table S2**. Primers for constructing pET23/42-*mlaCB* plasmid include MlaC-N5-NdeI (ATATCATATGTTTAAAC GTTTAATGATGGTCGCTTTGC) and MlaB-C3-XhoI (ATATCTCGAGTTAACGAGGCAGA ACATCAG). Luria-Bertani (LB) broth (Difco) was used for growing bacteria culture. Antibiotics (sigma) were added at the following concentrations, unless otherwise indicated: Ampicillin (Amp) was used at a concentration of 200 μg/mL for overnight (O/N) culture, 100 μg/mL for sub-culture; chloramphenicol (Cam) was used at 30 μg/mL; spectinomycin (Spec) was used at 25 μg/mL.

### Efficiency-of-plating

Cell strains were grown to OD_600_∼0.5-0.7 in LB with or without respective antibiotics with 1:100 dilution from an O/N culture at 37 °C with shaking. Then each strain was diluted to OD_600_∼0.1 and ten-fold serial-dilutions of each strain were prepared in LB in a 96-well microplate. 2 μL of culture from microplate was spotted to solid LB plates with or without adding SDS/EDTA or bile salts at the desired concentrations. Plates were incubated at 37 °C for O/N, and then the results can be visualized by G:BOX Chemi XT 4 (Genesys version 1.3.4.0, Syngene).

### Isothermal titration calorimetry (ITC)

ITC experiments were performed with Nano ITC (TA Instrument). Purified proteins (sMlaD and sLolB) were prepared at concentration of 500 μM, and bile salts were prepared at concentration of 4 mM (0.166%(w/v)) in PBS. In PBS, the reported CMCs of DOC and CDOC are ∼5.3 mM (0.220%) and ∼7 mM (0.290%), respectively (Simonović, 1997). DOC/CDOC was used at 4 mM to avoid significant heat changes due to micelle dissociations. ITC titration was set to 40 × 1.05 μl, 200s interval between injections, at 25 °C. Data were analyzed and fitted using the NanoAnalyze software (TA instrument) after subtracting background heat changes (from DOC-PBS titrations).

### In vitro photo-crosslinking with diazirine-DOC

16.5 μg of protein was mixed with 0.1% az-DOC (Santa Cruz Biotechnology) in PBS on a 96 well microplate, incubated for 5 minutes. Then, the mixture was (or not) subjected to ultraviolet (UV) light for 15 minutes. After UV treatment, the mixture was mixed with 2X Laemmli reducing buffer and boiled for 10 minutes at 100 °C, followed by SDS-PAGE and immunoblotting analyses. For competition, increasing concentrations of DOC/CDOC/cholate were added into the mixture before incubation.

## ACKNOWLEDGMENTS

This work was supported by the Singapore Ministry of Health National Medical Research Council under its Open Fund Individual Research Grant (MOH-000145) (to S.-S.C.). We thank J. Y. S. Chen and Department of Microbiology and Immunology, National University of Singapore for providing use of the ITC. We thank Taplin Mass Spectrometry Facility, Harvard Medical School for providing the tandem mass spectrometry service.

The authors declare no conflict of interest.

