## Supplementary information for "Deciphering the role of the OmpC-Mla system in bile salt resistance"

**Affiliations**:

**Supplementary Experimental**

**Protein expression and purification**

Affinity purification was performed according to a procedure described previously [1]. For BL21(λDE3) strains containing pET22/42 expressing MlaC/D, a 1.5-L culture was grown in LB broth with 1:100 dilution from an overnight culture adding respective antibiotics at 37 °C until $OD_{600}$~0.5–0.7. 1 mM IPTG (Axil Scientific, Singapore) was added and the culture was grown at 37 °C for another 3 h. Cells were collected by centrifugation at 4700 ×g for 20 min.

For holo-state proteins, 750 ml cell pellets were re-suspended in 20 ml 1X phosphate-buffered saline (PBS) buffer pH 7.4 containing 1 mM phenylmethylsulfonyl fluoride (sigma), 100 μg/ml lysozyme and 50 μg/ml DNase I in ice cold water. The re-suspended cells were lysed by a double passage through a high-pressure French press (French Press G-M, Glen Mills) homogenizer at 20000 psi. Lysed cultures were centrifuged at 4700 ×g for 10 min at 4 °C to remove unbroken cells. The cell lysate was collected and centrifuged at 100000 ×g for 1 h in an ultracentrifuge (Model XL-90, Beckman Coulter) at 4 °C. The supernatant was kept on ice. 4 ml of TALON cobalt resin (Clontech) was pre-equilibrated with 10 ml of washing buffer containing 1X PBS pH 7.4 at 4 °C for 30 minutes with rocking. The supernatant was loaded on the resin and incubated for 1h on ice while rocking. The resin mixture was later loaded onto the column and allowed to drain by gravity. The filtrate was collected, reloaded into the column, and drained as above. The column was washed with 2 × 10 ml washing buffer and 2 × 10 ml wash buffer adding 20 mM imidazole. The proteins were eluted with 8 ml of elution buffer which contains 1X PBS pH 7.4 and 200 mM imidazole. The eluate was concentrated in a 10 kDa cut-off ultra-filtration device (Amicon Ultra, Merck Millipore) by centrifugation at 4000 x *g* to 500~1000 μl.

For apo-state proteins, cell pellets were re-suspended in 20 ml of 1X PBS pH 7.4 and 8 M urea. The re-suspended cells were lysed with sonication on ice (20 % power, 1 second pulse on, 1 second pulse off for 5 mins). Lysed cultures were centrifuged at 4700 ×g for 10 min to remove unbroken cells. The cell lysate was collected and centrifuged at 100000 ×g for 1 h in an ultracentrifuge. The supernatant was kept. 4 ml of TALON cobalt resin was pre-equilibrated with 10 ml of 1X PBS pH 7.4 and 8 M urea for 30 minutes with rocking. The supernatant was loaded on the resin and incubated for 1h while rocking. The resin mixture was later loaded onto the column and allowed to drain by gravity. The filtrate was collected, reloaded into the column, and drained as above. The column was washed with 2 × 10 ml of 1X PBS pH 7.4 and 8 M urea adding 20 mM SDS and 2 × 10 ml of 1X PBS pH 7.4, 8 M urea, and 20 mM imidazole. The proteins were eluted with 8 ml of elution buffer which contains 1X PBS pH 7.4, 8 M urea, and 200 mM imidazole. The eluate was concentrated in a 10 kDa cut-off ultra-filtration device by centrifugation at 4000 xg to 500~1000 μl.

**SDS-PAGE and immunoblotting**

All samples subjected to SDS-PAGE were mixed with equal amounts of 2X Laemmli reducing buffer with or without boiling for 10 min at 100 °C. Equal volumes of the samples were loaded onto the gels. SDS-PAGE was performed according to Laemmli [2] using the 4% and 12% Tris·HCl stacking gels. After running SDS-PAGE, gels were either stained with coomassie brilliant blue staining (sigma) or subjected to immunoblotting.

Immunoblotting was performed by transferring protein bands from the gels onto polyvinylidene fluoride (PVDF) membranes (Immun-Blot® 0.2 μm, Bio-Rad) using semi-dry electroblotting system (Trans-Blot® TurboTM Transfer System, Bio-Rad). Transferred membranes were blocked by 1X casein blocking buffer (Sigma). α-DOC (Cloud-Clone Corp.) was used at 1:3000 dilution. α-rabbit IgG secondary antibody (GE Healthcare) conjugated to HRP (from donkey) was used at 1:5000 dilution. Luminata Forte Western HRP Substrate (Merck Milipore) was used to develop the membranes and protein bands were visualized by G:BOX Chemi XT 4.

**OM modification level measurement**

5 ml culture was grown in LB broth (with antibiotics if required) to $OD_{600}$ ~ 0.5-0.7 with adding in 1 μCi/ml $^{14}$C-acetate (PerkinElmer) from overnight (O/N) culture with 1:100 dilution at 37 °C. Cells were harvested by centrifuging at 4700 ×g for 10 minutes and the pellets were washed with 1 ml of 1X PBS buffer. Cells were re-suspended in 0.32 ml of 1X PBS buffer, 0.4 ml of chloroform (Fisher Chemical) and 0.8 ml of methanol (VWR) (single-phase Bligh-Dyer mixture, chloroform: methanol: water = 1: 2: 0.8) and incubated for 20 minutes with rocking at room temperature. The insoluble material was collected by centrifuging at 16000 ×g for 30 minutes. This insoluble material was washed with 1 ml of the single-phase Bligh-Dyer mixture followed by 16000 ×g centrifugation for collection and then re-suspended in 0.45 ml of 12.5 mM sodium acetate (sigma) pH 4.5 containing 1% SDS with 15 minutes sonication in a water bath sonicator. The re-suspended material was heated at 100 °C for 35 minutes. Next, 0.5 ml of chloroform and 0.5 ml of methanol were added into the mixture (forming the two-phase Bligh-Dyer mixture, chloroform: methanol: water = 2: 2: 0.8). The lower phase was collected by centrifuging for 10 minutes at 16000 ×g and washed with 1 ml of two-phase Bligh-Dyer mixture, collected by centrifuging for 10 minutes at 16000 ×g. The lower phase is dried O/N. Solid material lipid A was dissolved with 20 μl chloroform/ methanol (4:1) and radioactivity was measured using scintillation counter (Perkin Elmer MicroBeta2 scintillation). The thin layer chromatography (TLC) chamber is

prepared by adding in chloroform: pyridine (Merck): 88% formic acid (Merck): water (50: 50: 16: 5) and developed for O/N. The suspended lipid A with same radioactivity count was spotted onto TLC plate (Silica Gel 60 F254, Merck Millipore) and the plate is ran in the saturated chamber. The TLC plate was then dried and exposed to phosphor storage screens (GE Healthcare). Phosphor-screens were visualized in a phosphor-imager (Storm 860, GE Healthcare) and the spots were quantified by ImageQuant TL analysis software (version 7.0, GE Healthcare).

### Supplementary Tables

**Table S1. List of strains used in this study.**

| Strains | Reference |
|---|---|
| MC4100 [$F^-$ *araD139* Δ*(argF-lac) U169 rpsL150 relA1 flbB5301 ptsF25 deoC1 ptsF25 thi*] | [3] |
| MC4100 Δ*mlaA*::*kan* | [4] |
| MC4100 Δ*mlaC*::*kan* | [4] |
| MC4100 Δ*mlaD*::*kan* | [4] |
| MC4100 Δ*mlaF*::*kan* | [4] |
| MC4100 Δ*mlaE*::*kan* | [4] |
| MC4100 Δ*mlaB*::*kan* | [4] |
| MC4100 Δ*ompC*::*kan* | [4] |
| MC4100 Δ*ompC*::*ompC* | [5] |
| MC4100 Δ*ompC*::$ompC_{R92A}$ | [5] |
| MC4100 Δ*ompF*::*kan* | [4] |
| MC4100 Δ*ompC*Δ*ompF*::*kan* | [4] |
| MC4100 Δ*tolC*::*kan* | [4] |

**Table S2. List of plasmids used in this study.**

| **Plasmids** | **Plasmid Description**[a] | **Reference** |
|---|---|---|
| pET23/42 | pT7 inducible expression vector; Amp$^R$ | [6] |
| pET23/42-*mlaA-Chis* | Encodes full-length MlaA with C-terminal His$_8$ tag; Amp$^R$ (p-*mlaA*-His) | [4] |
| pET23/42-*mlaC-Chis* | Encodes full-length MlaC with C-terminal His$_8$ tag; Amp$^R$ (p-*mlaC*-His) | [4] |
| pET23/42-*mlaD-Chis* | Encodes full-length MlaD with C-terminal His$_8$ tag; Amp$^R$ (p-*mlaD*-His) | [4] |
| pET23/42-*mlaF-Chis* | Encodes full-length MlaF with C-terminal His$_8$ tag; Amp$^R$ (p-*mlaF*-His) | [4] |
| pET23/42-*mlaE-Chis* | Encodes full-length MlaE with C-terminal His$_8$ tag; Amp$^R$ (p-*mlaE*-His) | [4] |
| pET23/42-*mlaB-Chis* | Encodes full-length MlaB with C-terminal His$_8$ tag; Amp$^R$ (p-*mlaB*-His) | [4] |
| pET23/42-*mlaCB* | Encodes full-length MlaCB ; Amp$^R$ (p-*mlaCB*) | This study |
| pACYC184 | Low copy cloning vector; Cam$^R$ | [7] |
| pACYC184-*ompC-Chis* | Encodes full-length OmpC with C-terminal His$_8$ tag; Cam$^R$ (p-*ompC*-His) | [4] |
| pBR322 | Medium to high copy cloning vector; Amp$^R$ | [8] |
| pBR322-*pldA* | Encodes full-length OmpLA; Amp$^R$ (p-*pldA*) | [4] |
| pCDFDuet-$mlaB_{T52A}$-*Chis* | Encodes full-length $\text{MlaB}_{\text{T52A}}$ with C-terminal His$_8$ tag; Stp$^R$/Spt$^R$ (p-$mlaB_{T52A}$-His) | [1] |
| pCDFDuet-$mlaF_{K47L}$-*Chis* | Encodes full-length $\text{MlaF}_{\text{K47L}}$ with C-terminal His$_8$ tag; Stp$^R$/Spt$^R$ (p-$mlaF_{K47L}$-His) | [1] |
| pET22/42-s*mlaD-Chis* | Encodes soluble-domain MlaD with C-terminal His$_8$ tag; Amp$^R$ | [1] |
| pET22/42-*mlaC-Chis* | Encodes full-length MlaC with C-terminal His$_8$ tag; Amp$^R$ | [1] |

[a]*Amp$^R$: Ampicillin resistance; Cam$^R$: Chloramphenicol resistance; Stp$^R$: Streptomycin resistance; Spt$^R$: Spectinomycin resistance*

**Table S3: Summary of az-DOC-apo-sMlaD MS results.**

| ScanF | z | Peptide | ModScore Peptide | Position | Ascore[a] (0.3Da) | Ascore (0.6Da) |
|---|---|---|---|---|---|---|
| **11281** | 3 | R.T#EPTYTLYATFDNIGGLK.A | R.TEPTYT#LYATFDNIGGLK.A | 13 | 0.0 | 0.0 |
| **10815** | 2 | R.VAD#ITLDPK.T | R.V#ADITLDPK.T | 40 | 0.0 | 0.0 |
| **9863** | 3 | R.VADITLDPKTYL#PR.V | R.VADITLDPKTYL#PR.V | 51 | 0.0 | 0.0 |
| **10343** | 2 | R.VTL#EIEQR.Y | R.V#TLEIEQR.Y | 54 | 0.0 | 0.0 |
| **10432** | 2 | R.VTL#EIEQR.Y | R.VTL#EIEQR.Y | 56 | 31.3 | 42.9 |
| **7516** | 3 | R.YNHIPD#TSSLSIR.T | R.Y#NHIPDTSSLSIR.T | 62 | 0.0 | 0.0 |
| **7764** | 3 | R.YNHIPD#TSSLSIR.T | R.Y#NHIPDTSSLSIR.T | 62 | 0.0 | 0.0 |
| **7799** | 3 | R.YNHIPD#TSSLSIR.T | R.Y#NHIPDTSSLSIR.T | 62 | 0.0 | 0.0 |
| **10163** | 3 | R.YNHIPDTSSL#SIR.T | R.Y#NHIPDTSSLSIR.T | 62 | 0.0 | 0.0 |
| **10251** | 3 | R.YNHIPDTSSL#SIR.T | R.Y#NHIPDTSSLSIR.T | 62 | 0.0 | 0.0 |
| **7408** | 3 | R.YNHIPD#TSSLSIR.T | R.YNHIPD#TSSLSIR.T | 67 | 24.2 | 21.5 |
| **7631** | 3 | R.YNHIPD#TSSLSIR.T | R.YNHIPD#TSSLSIR.T | 67 | 24.2 | 30.3 |
| **7973** | 3 | R.YNHIPD#TSSLSIR.T | R.YNHIPD#TSSLSIR.T | 67 | 30.8 | 39.8 |
| **8159** | 3 | R.YNHIPD#TSSLSIR.T | R.YNHIPD#TSSLSIR.T | 67 | 24.2 | 30.3 |
| **8170** | 3 | R.YNHIPD#TSSLSIR.T | R.YNHIPD#TSSLSIR.T | 67 | 17.0 | 22.9 |
| **8282** | 3 | R.YNHIPD#TSSLSIR.T | R.YNHIPD#TSSLSIR.T | 67 | 17.5 | 21.5 |
| **8714** | 2 | K.D#GDTIQDTK.S | K.DGDTI#QDTK.S | 106 | 0.0 | 0.0 |
| **7308** | 4 | K.GD#DNKNSGDAPAAAPGNN ETTEPVGTTK.- | K.GD#DNKNSGDAPAAAPGNN ETTEPVGTTK.- | 129 | 5.7 | 6.7 |
| **6407** | 4 | K.GDDNKNSG#DAPAAAPGNN ETTEPVGTTK.- | K.GDDNKNSGD#APAAAPGNN ETTEPVGTTK.- | 136 | 0.0 | 0.0 |
| **7432** | 4 | K.GDDNKNSGDAPAAAPG#NN ETTEPVGTTK.- | K.GDDNKNSGDAPAAAPG#NN ETTEPVGTTK.- | 143 | 2.4 | 6.7 |

[a] Ambiguity score (Ascore). Ascore is the confidence score of assigning the position of crosslinked residues (#). Algorithm was run twice with fragment ion tolerance of 0.3Da and 0.6Da to avoid noise peak interference. The crosslinked position is considered confidently assigned (at 99% confidence level) if the Ascore values are both above 19 for the same site (highlighted in yellow) [9].

### Supplementary Figures

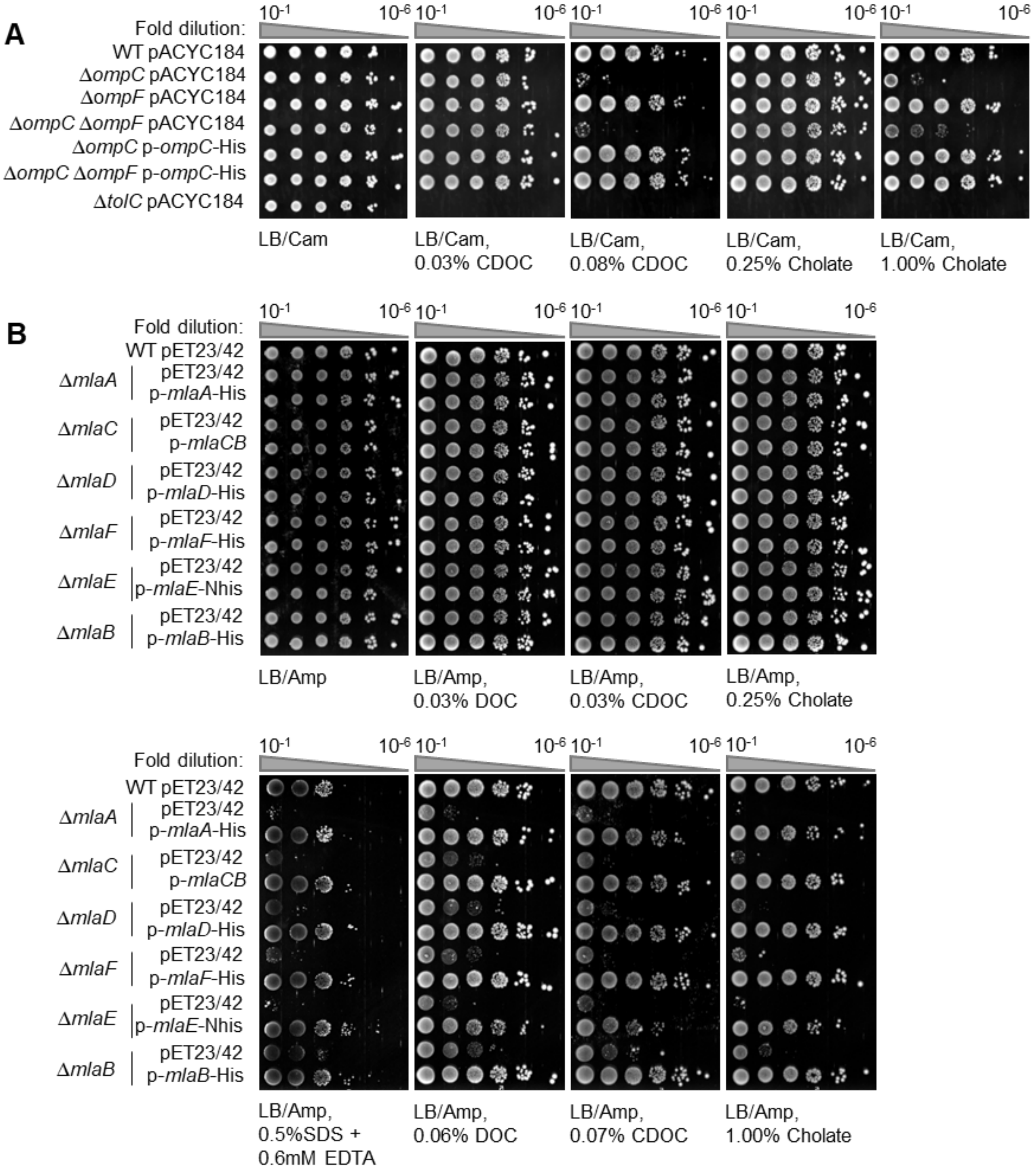


**Figure S1: Defects in the OmpC-Mla system confer cell bile salt sensitivity at/above CMCs.** EOP of indicated porin (A) or *mla* mutant (B) strains on LB media containing indicated concentrations of SDS/EDTA, DOC, CDOC or cholate. *mlaC* deletion has been shown to have effects on the expression of downstream genes *mlaB*/*yrbA* [9], and in this study MlaC and MlaB were expressed together to complement *mlaC* deletion.

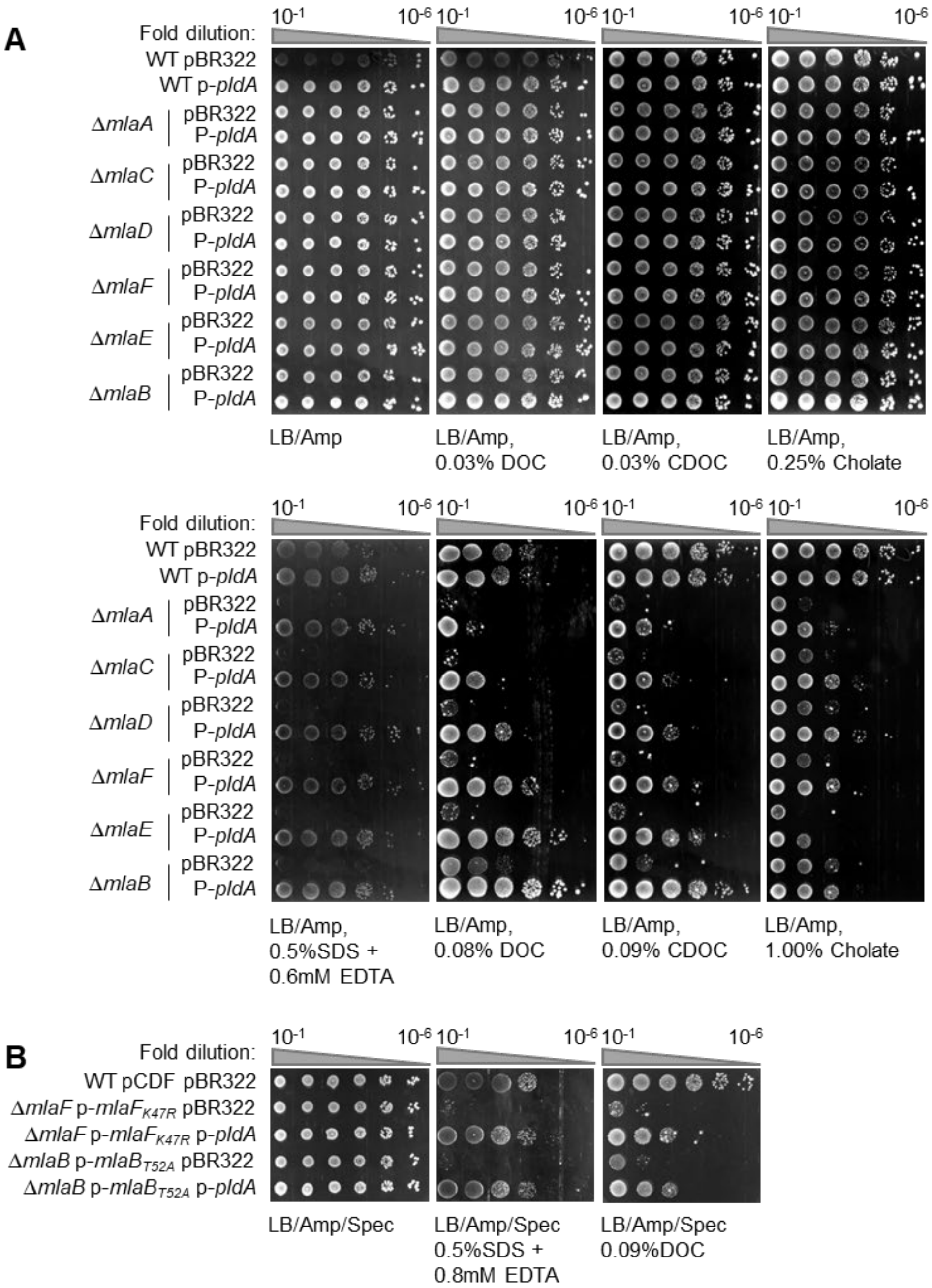


**Figure S2: Overexpression of PldA fully rescues SDS/EDTA sensitivity but only partially restores bile salt resistance in *mla* deletion and *mla* ATPase-defective mutants.** EOP of *mla* deletion (A) or ATPase-defective mutant (B) strains on LB media containing indicated concentrations of SDS/EDTA, DOC, CDOC, and cholate.

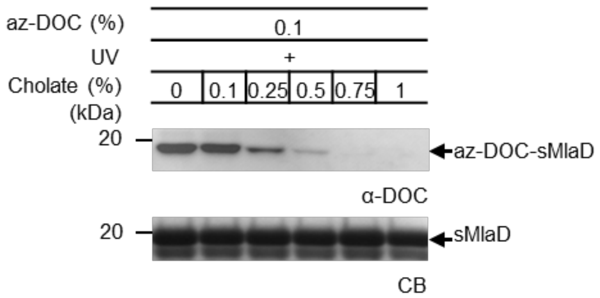


**Figure S3: Cholate can competitively prevent az-DOC crosslinking to soluble MlaD.** SDS-PAGE analysis and immunoblot showing UV-dependent formation of az-DOC-sMlaD adducts in the presence of increasing concentrations of cholate.

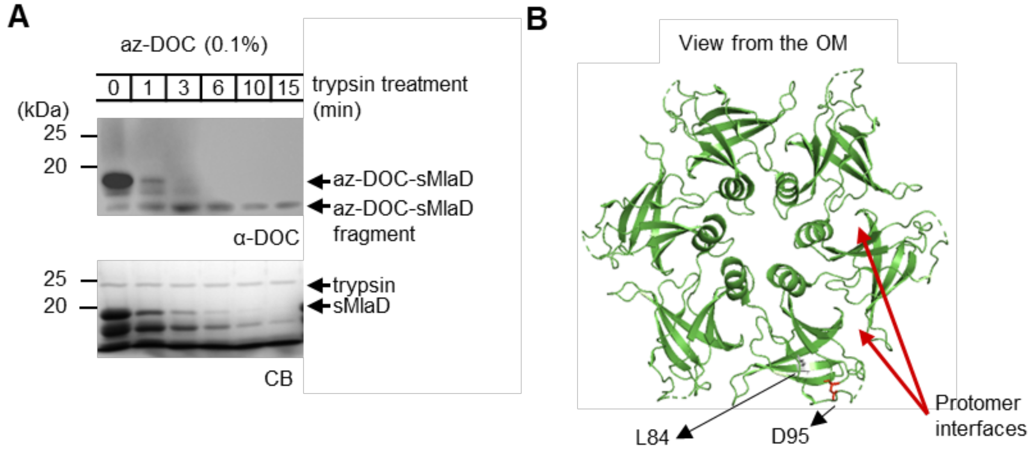


**Figure S4: Mass spectrometry of tryptic peptides derived from az-DOC-sMlaD crosslinked adducts reveal putative crosslinked sites close to MlaD protomer interfaces.** (A) SDS-PAGE analysis and immunoblot showing time-dependent trypsin digestion of az-DOC-sMlaD crosslinked adducts to give stable tryptic fragments. Total tryptic digest was subjected to LC-MS/MS to identify potential crosslinked residues (see **Table S3**). (B) Top view of cartoon representation of *E. coli* sMlaD (PDB: 5UW2) [10] with putative az-DOC-crosslinked residues inducated.

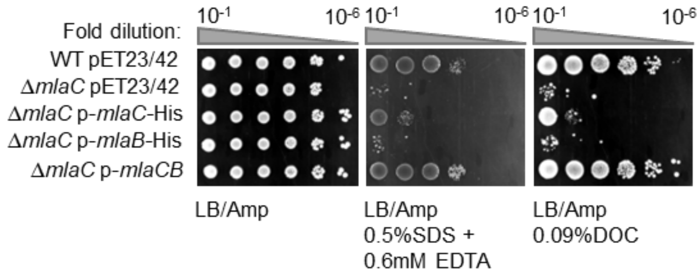


**Figure S5: MlaC is required for bile salt resistance.** EOP of *mlaC* mutant strains harboring empty vector (pET23/42) or plasmids expressing *mlaC*, *mlaB*, or *mlaCB*, on LB media containing indicated concentrations of SDS/EDTA and DOC.
